# Development of Web Application for the Comparison of Segment Variability with Sequence Evolution and Immunogenic Properties for Highly Variable Proteins: An Application to Viruses

**DOI:** 10.1101/2021.12.01.470810

**Authors:** Sumit Bala, Ambarnil Ghosh, Subhra Pradhan

## Abstract

High rate of mutation and structural flexibilities in viral proteins quickly make them resistant to the host immune system and existing antiviral strategies. For most of the pathogenic viruses, the key survival strategies lie in their ability to evolve rapidly through mutations that affects the protein structure and function. Along with the experimental research related to antiviral development, computational data mining also plays an important role in deciphering the molecular and genomic signatures of the viral adaptability. Uncovering conserved regions in viral proteins with diverse chemical and biological properties is an important area of research for developing antiviral therapeutics, though assigning those regions is not a trivial work. Advancement in protein structural information databases and repositories, made by experimental research accelerated the in-silico mining of the data to generate more integrative information. Despite of the huge effort on correlating the protein structural information with its sequence, it is still a challenge to defeat the high mutability and adaptability of the viral genomics structure. In this current study, the authors have developed a user-friendly web application interface that will allow users to study and visualize protein segment variabilities in viral proteins and may help to find antiviral strategies. The present work of web application development allows thorough mining of the surface properties and variabilities of viral proteins which in combination with immunogenicity and evolutionary properties make the visualization robust. In combination with previous research on 20-Dimensional Euclidian Geometry based sequence variability characterization algorithm, four other parameters has been considered for this platform: [1] predicted solvent accessibility information, [2] B-Cell epitopic potential, [3] T-Cell epitopic potential and [4] coevolving region of the viral protein. Uniqueness of this study lies in the fact that a protein sequence stretch is being characterized rather than single residue-based information, which helps to compare properties of protein segments with variability. In current work, as an example, beside presenting the web application platform, five proteins of SARS-CoV2 was presented with keeping focus on protein-S. Current web-application database contains 29 proteins from 7 viruses including a GitHub repository of the raw data used in this study. The web application is up and running in the following address: http://www.protsegvar.com.

## 1. Introduction

Pandemic situation is an outcome of the newly emerged pathogenic strain where majority of the host population does not have any immunity against the pathogen and as a result high mortality and morbidity are observed over the affected areas. Coronavirus 2020 pandemic is the most recent example which includes the examples of both viral mutability and host’s immune resistance against it. Coronavirus infection was first reported in chicken in the late 1920s (Estola, 1970) and human coronaviruses were discovered in the 1960s (Kahn and McIntosh, 2005). B814 was the name assigned to the first coronavirus strain and currently it is not clear that which strain represents that origin (Corman et al., 2014). Before the current SARS-CoV-2, infective human coronavirus strains appeared several times. It was SARS-CoV in 2003, HCoV NL63 in 2003, MERS-CoV in 2013 and few more strains. Though they belong to the same virus group but each time they appeared with mutations which are novel and infective to human immune systems. Many animal coronaviruses were identified since 1960 and till now it’s a quite convincing theory that the pathogenic strains in coronaviruses have high similarities and emerged from bat and avian origins (Zhou et al., 2020, Wu et al., 2020, Mollentze and Streicker, 2020). Like coronavirus, influenza is another recurring pandemic causing virus and emerged several times with small variations in their genome. Influenza pandemic that infamous by the name of 1918 Spanish flu killed ∼20 to 50 million people worldwide. Followed by its Asian flu in 1957, Hongkong flu in ∼1968, Russian flu in 1977, H5N1 bird flu in 2008 and swine flu in 2009 left their footprint in human immunity with death and survival of the hosts. Small-pox, Yellow-fever, measles, etc. are among other pandemic viruses.

One of the major reasons for recurring pandemic events and high level of infectivity of a virus is mutation. Evolution of viral genome is a unique combination of various mechanisms like high mutation rate, segment reassortment, host shifting, improving virion stability, significant changes to their protein structure, etc. and more. Though their unit structures are very simple, but still it is really a challenge for viral system to hold functional integrity of their proteins tolerating the mutations accumulated in a random manner on their protein (Peck and Lauring, 2018). Novel mutations are the main reason behind inactivation of existing antiviral strategies. Known inhibitors of influenza viruses are amantadine, rimantadine, oseltamivir and zanamivir and currently resistance have developed in almost all circulating strains of flu for the first two drugs. H274Y mutation of neuraminidase protein is responsible for an oseltamivir resistance mutant, which is positioned in the active site of the viral protein. Q136K is another example of resistant mutant which is believed to responsible for zanamivir resistance. HIV is a rapidly mutating virus; nucleoside analog azidothymidine (AZT) was the first approved drug against it and rapid nature of the mutation lead to resistance (Boyer et al., 2006). A different strategy often gets highlighted in viral infection when mutations help the virus to evade host immune system. A recent study on Zikavirus NS1 protein it is shown that A188V mutation on the same protein helps the virus to evade immune response and potentiate infection (Xia et al., 2018). This is true for HBV also, showing a series of point mutations associated with immune escape and also resulting in failure in vaccination (Coppola et al., 2015). Therefore, recurrence of novel mutated viral strains is the reason behind frequent change in vaccination regimen. A well-known example is influenza virus which constantly evolve and bring changes to their molecular structure leads to yearly update vaccine candidates (Goodwin et al., 2006). Continuous accumulation of mutations on viral genome hardly indicates the error of polymerase, rather it comes from the absence or fault of proofreading mechanisms (Sanjuán and Domingo-Calap, 2016). Finally, not always mutations in the pandemic causing strain affects the pandemic situation, but for the emergence of new strain and drug resistance mutations are the key player. Therefore, it is very important to study mutations and their occurrences with the evolution of viral genetic material.

Multiple Sequence Alignment (MSA) based visualization and resulting sequence variability profile extraction is a widely used choice for the study of mutations in highly similar aligned sequences collected from different strains of viruses (Edgar and Batzoglou, 2006). As an example, PVS is a MSA based sequence variability analysis software which explores the conserved 3D structure and epitopic potentials of viral proteins (Garcia-Boronat et al., 2008). HotSpot3D is another recent server which analyzes mutations on a protein 3D structure and this server based on a software that is widely used for similar purpose (Chen et al., 2020). In 2009, one of the authors of the current manuscript published a 20D Euclidian space based algorithm for protein sequence and variability characterization including a phylogenetic distance matrix generator (Nandy et al., 2009). The algorithm successfully converted the variation in a protein sequence to qualifiable numerical values and later it was applied to detect variability in the drug targetable proteins of influenza and rotavirus (Ghosh et al., 2012, GHOSH and NANDY, 2011, Ghosh et al., 2010, Ghosh et al., 2009). On neuraminidase proteins of influenza-A virus (H5N1) this algorithm detected size regions which can be targeted for combating the virus. Indeed, after 3D mapping on crustal structure it was found that four of the regions covered by patches on active site of the protein (Ghosh et al., 2010). Later similar method was applied on VP7 protein of rotavirus and four such surface exposed conserved regions were found (Ghosh et al., 2012).

In this current work, a comprehensive protein stretch variability visualizing platform (web-application) is developed and applied to viral proteins. This approach standout from others on the basis that it is a stretch variability miner than a single position of amino acids. In addition, the graphical interface has scope that will allow users to browse through different stretch length to ensure its comparability with other length based structural and immunological features of the proteins. Currently, the plot compares stretch variability with four important properties of protein stretch (peptides): solvent accessibility, B-Cell epitope, T-Cell epitope and coevolution within the residues. Addition of coevolution residue pairs maps to the variability helps the users to prioritize the conserved stretches with coevolution hotspots. Beside getting idea on coevolution profile, the two most common application of protein stretch mining is the in-silico selection of vaccine candidates and screening of important surface regions from proteins. Important surface regions not only help to identify proteins’ active site and allosteric sites, but also may help to find unknow functions of proteins and vulnerability spots for drug targeting.

## 2. Materials and methods

### 2.1. Sequence Collection, Processing and coevolution

Protein sequences analyzed in the current study were collected from NCBI Virus, *sequence for discovery* portal (Hatcher et al., 2017) (https://www.ncbi.nlm.nih.gov/labs/virus/vssi/#/). Total 7 pandemic/endemic human disease-causing viruses were selected according to the list in World Health Organization (https://www.who.int/emergencies/diseases/en/) and proteins were chosen based on the environmentally exposed part of the virions. For the processing and preparation of FASTA file SeqKit FASTA/Q file manipulation tool were used (Shen et al., 2016) (https://github.com/shenwei356/seqkit) and multiple alignment were performed in MUSCLE (Edgar, 2004) (http://www.drive5.com/muscle/). pyDCA package were used for the purpose of coevolution analysis (Zerihun et al., 2020) (https://pypi.org/project/pydca/).

### 2.2. Calculation of 20D algorithm-based variability of proteins

Variability calculation of the viral protein sequence stretches were performed using 20-dimensional Cartesian coordinate based graphical representation algorithm (Nandy et al., 2009, Ghosh et al., 2010). The variability detection pipeline used in this work was previously studied by Ghosh et al. on surface proteins of influenza and rotavirus (Ghosh et al., 2012, Ghosh et al., 2010). In brief, this algorithm compares sequences in a 20D space by associating each amino acid with a single axis of twenty. Degeneracy in the output for dissimilar protein stretches were corrected by using weighted average, resultant vector and with addition of an initial weightage to each coordinate. This algorithm also performed phylogenetic analyses for a bunch of protein in 20D space (Nandy et al., 2009), which functionality was not included in the work-flow.

### 2.3. Prediction of solvent accessibility & immunological potential of peptides

For the purpose of solvent accessibility and immunological property prediction, reference sequences were considered as standard for each viral protein; as an example in SARS-CoV2 coronavirus strain isolate Wuhan-Hu-1 was (https://www.ncbi.nlm.nih.gov/nuccore/NC_045512.2) considered for this purpose. Residue wise solvent accessibility of the amino acid residues were predicted using standalone NetSurfP application version 2.0 (Klausen et al., 2019) (http://www.cbs.dtu.dk/services/NetSurfP/) and average solvent accessibility of the stretches were made according to the window length along the protein sequence. Similarly T-Cell and B-Cell epitopic properties were predicted using standalone version of IEDB Epitope Prediction and Analysis Tools (Vita et al., 2019) (http://tools.iedb.org/main/).

### 2.4. Development of web application

Web application was developed in Flask microframework, with a front-end user interface (UI) designed in JavaScript and backend in Python. In **Figure 1**, a schematic diagram explaining the server was presented with details of the data processing directions. This web application was designed to perform two types of job: [a] A simple pR-value calculator, where user can submit a single protein sequence to obtain pR-value (Ghosh et al., 2010) (**Figure 1**). [b] the second one is the main goal the server is designed for, which is a stretch variability explorer where from the drop-down menu user will be able to select the virus name and specific protein to get the comparative graphical presentation within the output box (marked No. 4, **Figure 2**).

**Figure 1:**
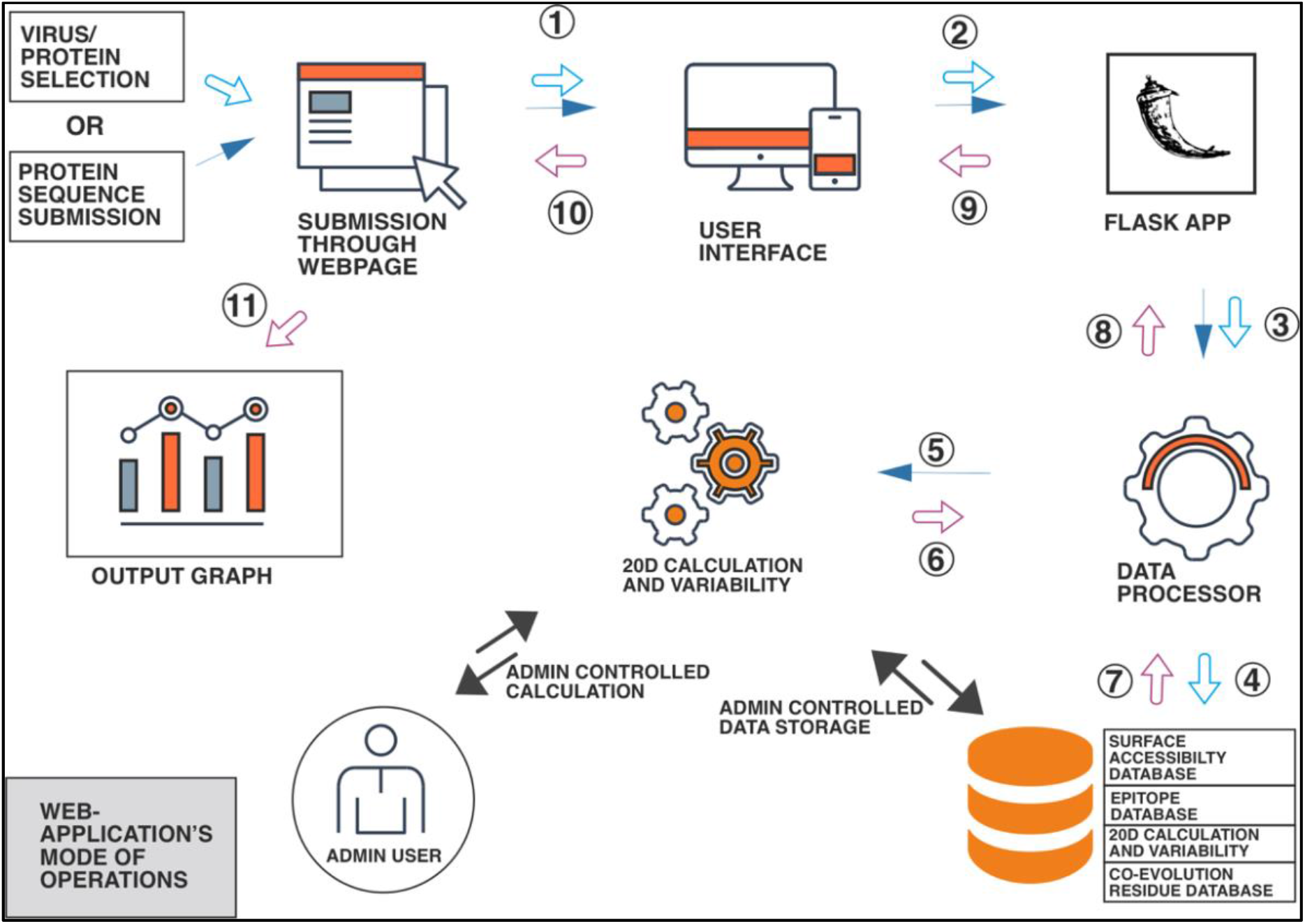
Schematic diagram explaining the workflow of the server. Inputs were indicated in blue arrow (steps 1, 2, 3, 4 and 5) and outputs follow the red arrows (steps 6, 7, 8, 9, 10 and 11). Black arrow only indicates the internal operation of the server between installed code and databases. Admin user in this interface is responsible computations and database updates.

**Figure 2:**
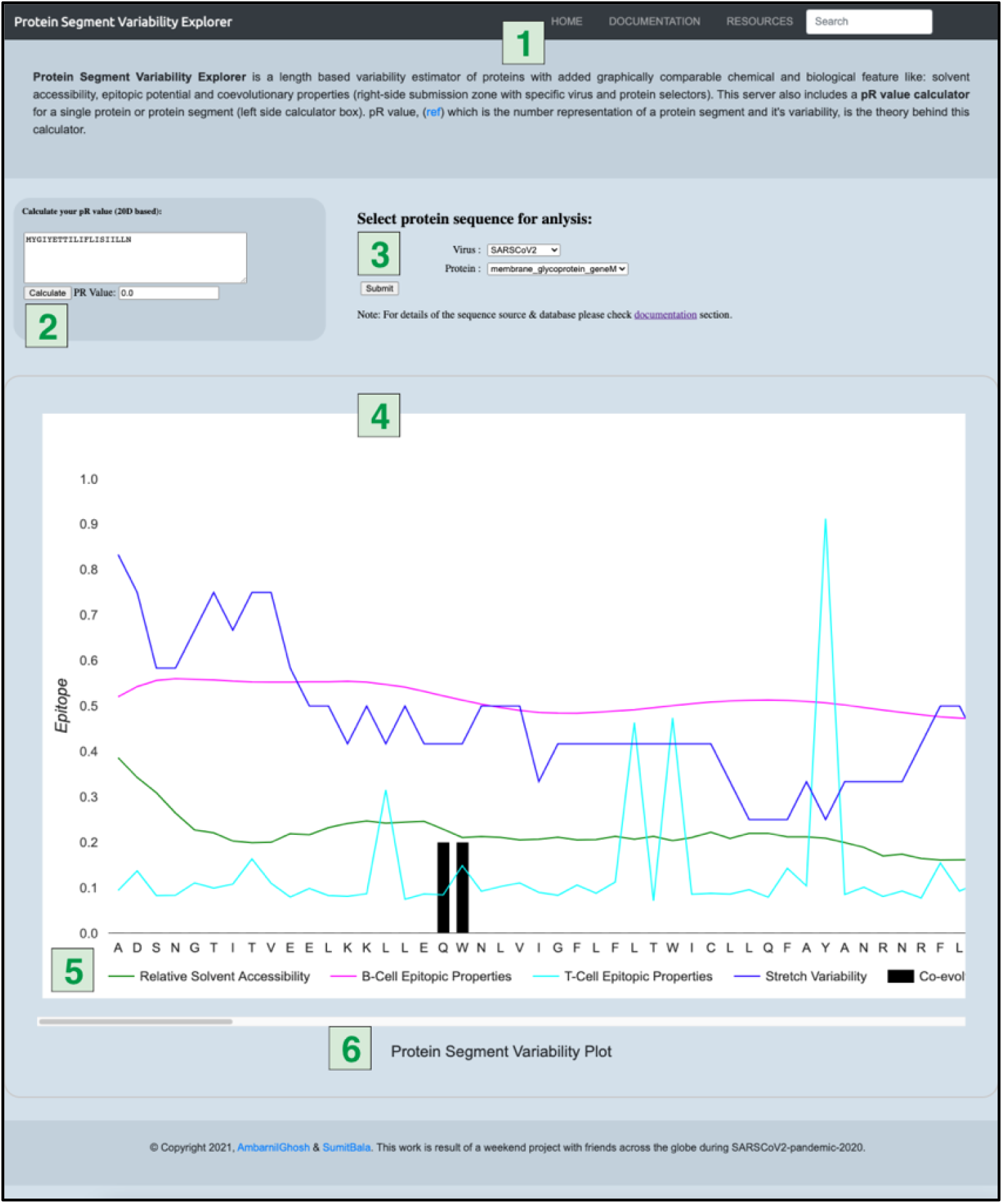
Explanation of the user interface from the front page of the server. Marked No.1 indicates the menu bar where documentation and references can be browsed. Both No. 2 and 3 are submission form where No.4 showed the output plot area. No. 5 and 6 were shown to indicate the axes details and scroll-bar necessary for browsing through plot’s content.

The submission form of the web application platform was designed in HTML and available to users in through website http://www.protsegvar.com. As mentioned previously the interface has two submission forms incorporated in a single web page (mentioned in No. 3 and 4 in **Figure 2**). As the output graph compares data related to multiple properties of a protein segment, epitope and solvent accessibility scores are presented in X-axis scale and keeping their original values as obtained from the prediction program. In case of T-Cell epitope value a higher threshold cut-off of 1 is programmed indicating a high epitopic potential when the peak crosses that threshold.

Though Y-Axis indicates the position of the segment of length 9 amino acids, where each of the single letters indicates the starting amino acid of that 9 amino acid long segment. Variability was shown in blue line and number of unique sequences obtained for that segment was indicated in right-Y axis which can only be seen at the end of the sequence. As the co-evolving residue pairs comes in a score-based priority, authors decided to provide only top 10 residues in this plot. The rest of the coevolving residues will be available from the above-mentioned GitHub repository created for this server’s data (**Section 2.5**). When dragging through the horizontal scroll bar (showed in No. 6 of **Figure 2**), last 8 positions of each plot come with zero values because the plot only consider 9 amino acid segments.

### 2.5: Redistributable Data Repository in GitHub

The raw data used for rendering the graphical interface is available through in form a database uploaded in GitHub repository https://github.com/ambarnilghosh/psvarDB-20D. Currently the repository contains the database of 7 pandemic/endemic viruses with their 29 proteins. The database contains five parameters for each of the proteins considered to render in this web interface: [1] stretch variability data, [2] predicted B-Cell variability data, [3] predicted T-Cell variability data, [4] predicted surface accessibility data and [5] pairwise top20 coevolving residue information in separate files. Data updated according to the new proteins’ candidates will be updated to the server and uploaded in the GitHub repository.

## 3 Result & Discussion

Protein segment variability plot presented in the current web-application not only include the variability of a region of certain length but also provide useful information where this segment specific variability can be compared with four important biological information related to surface accessibility, co-evolution and immunological properties. Though in the initial version of the server it stored and represented the results of proteins from seven viruses (included in coronavirus and flu group), in this section only the spike protein of SARS-CoV2 was presented and explored as an example of this server’s application. In addition, also other four proteins (protein E, protein M, ORF3a and ORF8) were discussed for their coevolving residues and Sequence of Interests (SOI). SOI regions can be defined as the segment of protein where graph showed low variability, more surface exposure, and immunologically potential regions. Though complete and high resolution PDB structures are not present for all the proteins, but in case of our protein-S example PDB structure is present (PDB id 7DDD) & the results are explained through that structure (Figure 4).

After running the aligned protein sequences of SARS-CoV-2 Protein S through our web application the resulting data were plotted in **Figure 3** after rescaling adjustments, avaeraging and normalization. A window size of 9 amino acid were considered for this analysis to produce the graph as a sample segment size (**Figure 3)**. Beside the variability and immunological properties top 20 scoring coevolving residues pairs are showed as a black dot marker closest to X-axis. Though the presentation does not explain about the distance between coevolving residues, it represents the position of the residues which were involved in the coevolving phenomena. From the **Table 1** and Protein S column, it’s clear that all the top20 coevolving comes is contagious positions. But for protein ORF8 and protein ORF3a interesting residue couples were found, which were discussed in the next paragraphs. From the pattern of the graph, author chose nine sample regions containing SOI pointing to the conserved stretches on the spike protein (SOIs were shown in orange bar in **Figure 3**). Choice of SOI will completely depend on the user i.e., according to their requirements on the proteins variability and other properties they will be able to choose the regions. As an example, regions with lowest variability (conserved sequences) and also with the highest surface accessibility and immunogenic properties can extract good epitope candidates from the graph which may help in designing vaccine candidates (Ghosh et al., 2010, Ghosh et al., 2011).

**Table 1:** Tabular description of the top 20 coevolving residue copuple in the five proteins of SARS-CoV-2.

| Top20 residues | protein S | protein E | protein M | protein ORF3a | protein ORF8 |
| --- | --- | --- | --- | --- | --- |
| 1 | 258-259 | 43-44 | 19-20 | **196-224** | 91-92 |
| 2 | 255-256 | 23-24 | 212-213 | 272-273 | 64-65 |
| 3 | 760-761 | 72-73 | 210-211 | 176-177 | 85-86 |
| 4 | 256-257 | 56-57 | 129-130 | 189-190 | 79-80 |
| 5 | 319-320 | 74-75 | 87-88 | 162-163 | 3-6 |
| 6 | 260-261 | 69-70 | 191-192 | 186-187 | 13-14 |
| 7 | 840-841 | 53-55 | 114-115 | 93-94 | 34-35 |
| 8 | 241-242 | 17-19 | 21-22 | 160-161 | 4-5 |
| 9 | 238-239 | 58-59 | 115-116 | 87-88 | 86-89 |
| 10 | 544-545 | 17-18 | 127-128 | 163-164 | **34-89** |
| 11 | 255-257 | 16-18 | 104-105 | 159-161 | 85-89 |
| 12 | 628-629 | 65-66 | 207-208 | 101-102 | 31-32 |
| 13 | 418-419 | 61-62 | 213-214 | 85-86 | 37-38 |
| 14 | 158-159 | 36-37 | 195-196 | 174-175 | **34-86** |
| 15 | 261-262 | 35-37 | 101-102 | 191-192 | **34-85** |
| 16 | 522-523 | 67-68 | 16-17 | 121-122 | 9-11 |
| 17 | 546-547 | 59-60 | 186-188 | 120-121 | 91-93 |
| 18 | 446-447 | 35-36 | 194-195 | 157-158 | 25-26 |
| 19 | 967-968 | 29-31 | 187-188 | 162-164 | **6-87** |
| 20 | 242-243 | 16-17 | 124-125 | 140-141 | 9-10 |

**Figure 3:**
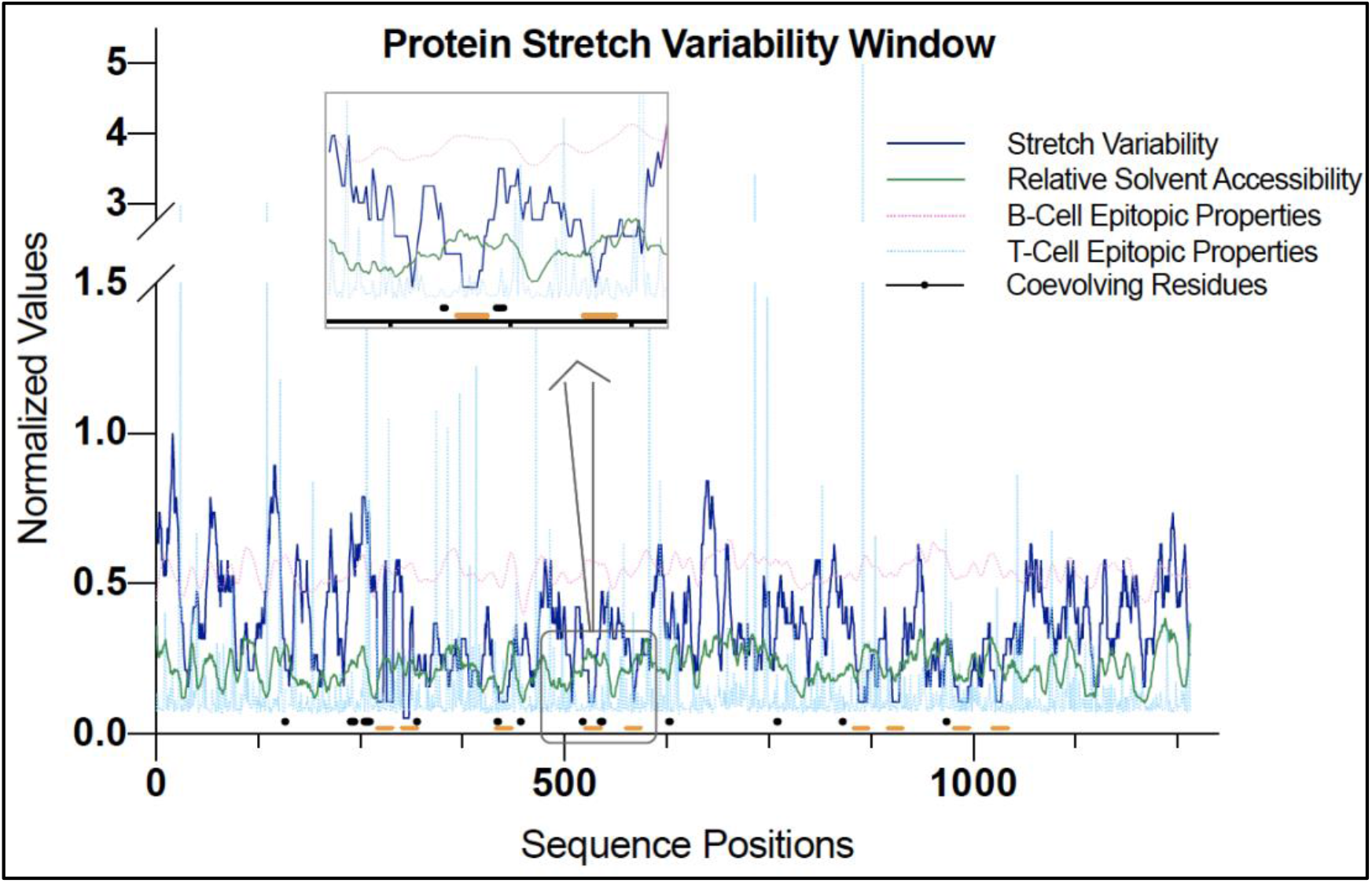
Presentation of protein stretch variability in a comparative graphical illustration for the spike protein (protein S) of SARS-CoV2. The color coding is like what presented in web-application platform.

From the candidate Protein-S, among 9 SOI regions, the region around amino acid number 540 shows strong conserved stretch with high B-cell epitopic activity. Another case is the region around amino acid number 880 showed all four properties perfectly favorable for an interesting stretch where sequence conserveness is considerable, solvent accessibility is high and both B and T-cell epitopic properties were also prominent. Authors presented all nine peptides within the 3D structure of Spike protein in **Figure 4** where targeted regions are presented in fluorescent green color. In the current version, the web application does not show the SOI by itself and it is expected that researchers will be able to dig their SOI according to requirements. In a future version of this server authors have plan to incorporate the SOI identification tool with the graph.

**Figure 4:**
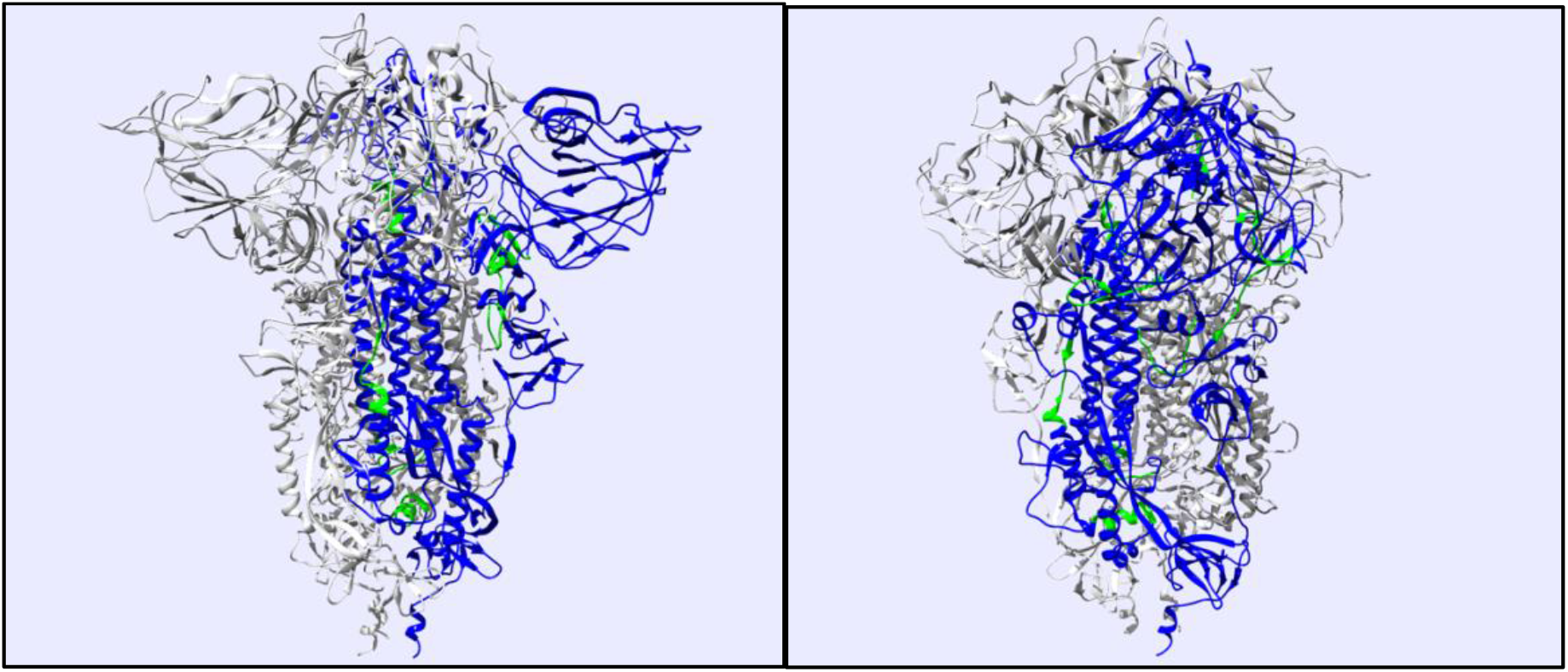
Presentation of nine SOI in the 3D structure of SARS-CoV-2 protein S. The SOIs are marked with fluorescent green color on the blue colored protein chain A.

Through the study of coevolving residues interesting information emerged in the proteins which are considered for the current sample set. With the first four parameters, co-evolving residues add a weightage to the region under consideration because of the presence of a good coevolution score explain the higher variability. Highly variable regions with the pressure of coevolving residue not only explain conserveness of those regions, but also they indicate the key player in protein structure formation. Specifically, in viral molecular biology, presence of coevolving residues which are placed apart from each other have immense importance (Champeimont et al., 2016). Here in protein S, none of the high scoring coevolving pair showed distance among them following the similar result for protein M and protein E. But, in ORF3a and ORF8, interesting distances of coevolving residues are found (marked as bold letters in **Table 1**). Top scoring couple in ORF3a (residue no. 196 and 224) showed a distance of ∼28 amnio acids. In protein ORF8, two regions around amino acid number 6 and 34 showed good coevolution score with reside around 85 to 89.

Like the proteins of SARS-CoV-2 viruses presented above, we have considered six other viruses considering all total 29 proteins: H1N1 flu virus, H2N2 asian flu virus, H3N2 HongKong flu virus, H5N1 bird flu virus, MERS coronavirus and SARS coronavirus. We are currently working on updating the server database with more viruses and application features. Currently the server only runs with a window of nine amino acid stretch, but a development on the stretch length selection method is underway and feature will be deployed soon with further development in the interface.

## 5. Acknowledgement

This work was performed as a weekend project and the resources consumed for this work was provided personally by authors. There is no potential competing interest is involved with this work with any of the Authors.

## 6. Author’s Contribution

SB and AG were involved in the server development where SB did the web application development, where AG helped him with data organization and webpage design. Co-evolution concept added to this algorithm by SP & AG; and the final implementation plan was designed by AG and SB. Manuscript was written by AG, SB & SP.

## 7. Supplementary Data

Supplementary data related to coevolving residues and other results for all 29 proteins can be found in GitHub repository: https://github.com/ambarnilghosh/psvarDB-20D

